# A multi-phase multi-objective dynamic genome-scale model shows different redox balancing among yeast species in fermentation

**DOI:** 10.1101/2021.03.08.434513

**Authors:** David Henriques, Romain Minebois, Sebastian Mendoza, Laura G. Macías, Roberto Pérez-Torrado, Eladio Barrio, Bas Teusink, Amparo Querol, Eva Balsa-Canto

## Abstract

Yeasts constitute over 1500 species with great potential for biotechnology. Still, the yeast *Saccharomyces cerevisiae* dominates industrial applications and many alternative physiological capabilities of lesser-known yeasts are not being fully exploited. While comparative genomics receives substantial attention, little is known about yeasts’ metabolic specificity in batch cultures. Here we propose a multi-phase multi-objective dynamic genome-scale model of yeast batch cultures that describes the uptake of carbon and nitrogen sources and the production of primary and secondary metabolites. The model integrates a specific metabolic reconstruction, based on the consensus Yeast8, and a kinetic model describing the time-varying culture environment. Besides, we proposed a multi-phase multi-objective flux balance analysis to compute the dynamics of intracellular fluxes. We then compared the metabolism of *S. cerevisiae* and *S. uvarum* strains in a rich medium fermentation. The model successfully explained the experimental data and brought novel insights into how cryotolerant strains achieve redox balance. The proposed model (along with the corresponding code) provides a comprehensive picture of the main steps occurring inside the cell during batch cultures and offers a systematic approach to prospect or metabolically engineering novel yeast cell factories.

**IMPORTANCE:** Non-conventional yeast species hold the promise to provide novel metabolic routes to produce industrially relevant compounds and tolerate specific stressors, such as cold temperatures. This work presented and validated the first multi-phase multi-objective genome-scale dynamic model to describe carbon and nitrogen metabolism throughout batch fermentation. To test and illustrate its performance, we considered the comparative metabolism of three yeast strains of the Saccharomyces genus in rich medium fermentation. The study revealed that cryotolerant Saccharomyces species might use the GABA shunt and the production of reducing equivalents as alternative routes to achieve redox balance, a novel biological insight worth being explored further. The proposed model (along with the provided code) can be applied to a wide range of batch processes started with different yeast species and media, offering a systematic and rational approach to prospect non-conventional yeast species metabolism and engineering novel cell factories.

## INTRODUCTION

Yeasts have been used to produce fermented foods and beverages for millennia and are among the most frequently used microorganisms in biotechnology. *Saccharomyces cerevisiae* dominates the scene and many research efforts focus on engineering this species for particular applications (e.g., (1, 2, 3) or (4)). Nowadays it is used to produce glycerol (5), biopharmaceutical proteins (6), or secondary metabolites, such as aroma our bioflavours (7, 8).

However, yeasts constitute a large group of 1500 (so far) described species and much less attention has been paid to non-conventional yeasts. These species remain a mostly untapped resource of alternative metabolic routes for substrate use and product formation as well as tolerances to specific stressors (9, 10). To exploit these alternatives efficiently, it is essential to understand the metabolic pathways of these species. Given the complexity of the endeavor, a modeling approach becomes indispensable.

Genome-scale models (GEMs) can contextualise high-throughput data and predict genotype-environment-phenotype relationships (11, 12). While GEMs have been widely used for the study and metabolic engineering of *S. cerevisiae* strains in continuous (steady-state) fermentations (13), their use to predict batch (dynamic) fermentation is still scarce. Nevertheless, many yeast-based processes operate in batch mode.

In batch operation, cell culture follows a growth curve with the following phases: lag-phase, exponential growth, growth under nutrient limitation, stationary phase, and cellular decay. Available dynamic GEMs of yeast metabolism focus on the exponential phase and explain reasonably well the measured dynamics of biomass growth, carbon sources uptake, and the production of relevant primary metabolites (14, 15, 16, 17). The development of GEMs that describe the five phases of batch processes, considering carbon and nitrogen metabolism throughout time and explaining secondary metabolism, is still required.

In this work, we derived a multi-phase and multi-objective dynamic genome-scale model of batch fermentation, which accounts for carbon and nitrogen metabolism throughout time and explains secondary metabolism. The model required various refinements to succeed: i) a novel metabolic reconstruction, based on an extension of the current consensus genome-scale model of *S. cerevisiae* (Yeast8, (18)); ii) multi-phase multi-objective implementation of a parsimonious flux balance analysis (pFBA, (19)) to compute the dynamics of the intracellular fluxes; iii) a model of protein turnover to explain nitrogen homeostasis and iv) a dynamic biomass equation to account for biomass composition variations throughout the process.

As relevant case study we considered the metabolism of *S. cerevisiae* and *S. uvarum* strains in rich medium fermentation. Recent studies revealed that *S. uvarum* strains show interesting physiological properties. *S. uvarum* is more cryotolerant than *S. cerevisiae,* produces more glycerol and less ethanol than *S. cervisiae* wine strains, and different aroma profiles (20, 21, 22, 23). In addition, traits such as its increased 2-phenylethanol (24, 25) yield could make this species a good candidate for metabolic engineering studies (26).

We applied the proposed model to investigate the origin of the phenotypic divergence between species. The model explained the experimental data successfully and revealed differences into how species achieve redox balance. Predicted intracellular fluxes led us to hypothesize that cryotolerant yeast strains can use the GABA shunt as an alternative NADPH source and store reductive power-necessary to subdue oxidative stress under cold conditions-in lipids or other polymers. Additionally, our results are compatible with recent experimental observations showing that most carbon skeletons used to form higher alcohols (i.e., isoamyl alcohol, isobutanol, and 2-phenylethanol) are synthesized *de novo.*

## RESULTS

### The novel metabolic reconstruction

We updated the Yeast8 consensus genomescale reconstruction of *S. cerevisiae* S288C (v.8.3.1) (18) to include 38 metabolites and 50 reactions to explain secondary metabolism (Table S1). Furthermore, comprehensive metabolic annotations, such as BO terms and MetaNetX identifiers, were added to the new metabolites and reactions.

Among the metabolites added, 13 aroma compounds were included. Noticeably, we found that prior genome-scale reconstructions lacked methionol and tyrosol impeding simulated growth on methionine and tyrosine as sole nitrogen sources, which is known to be possible for several *S. cerevisiae* strains, including S288C.

MetaDraft, AuReMe, and the results from the orthology analysis were used to create strain-specific models for two wine strains *S. cerevisiae* T73 and *S. uvarum* BMV58 and a strain *S. uvarum* CECT12600 found in non-fermentation environments (further details can be found in Text S1). Strains will be denoted as ScT73, SuBMV58 and SuCECT12600 from now on. The three models had 2, 3 and 2 reactions that were not in Yeast8, respectively.

### The multi-phase multi-objective flux balance analysis framework

Our results showed that batch fermentation modeling should be divided into five phases in which cellular objectives and flux constraints need to be modified: lag phase, exponential growth, growth under nitrogen limitation, stationary, and decay (Figures 1.A-1.B sketch the modeling approach).

**FIG 1.**
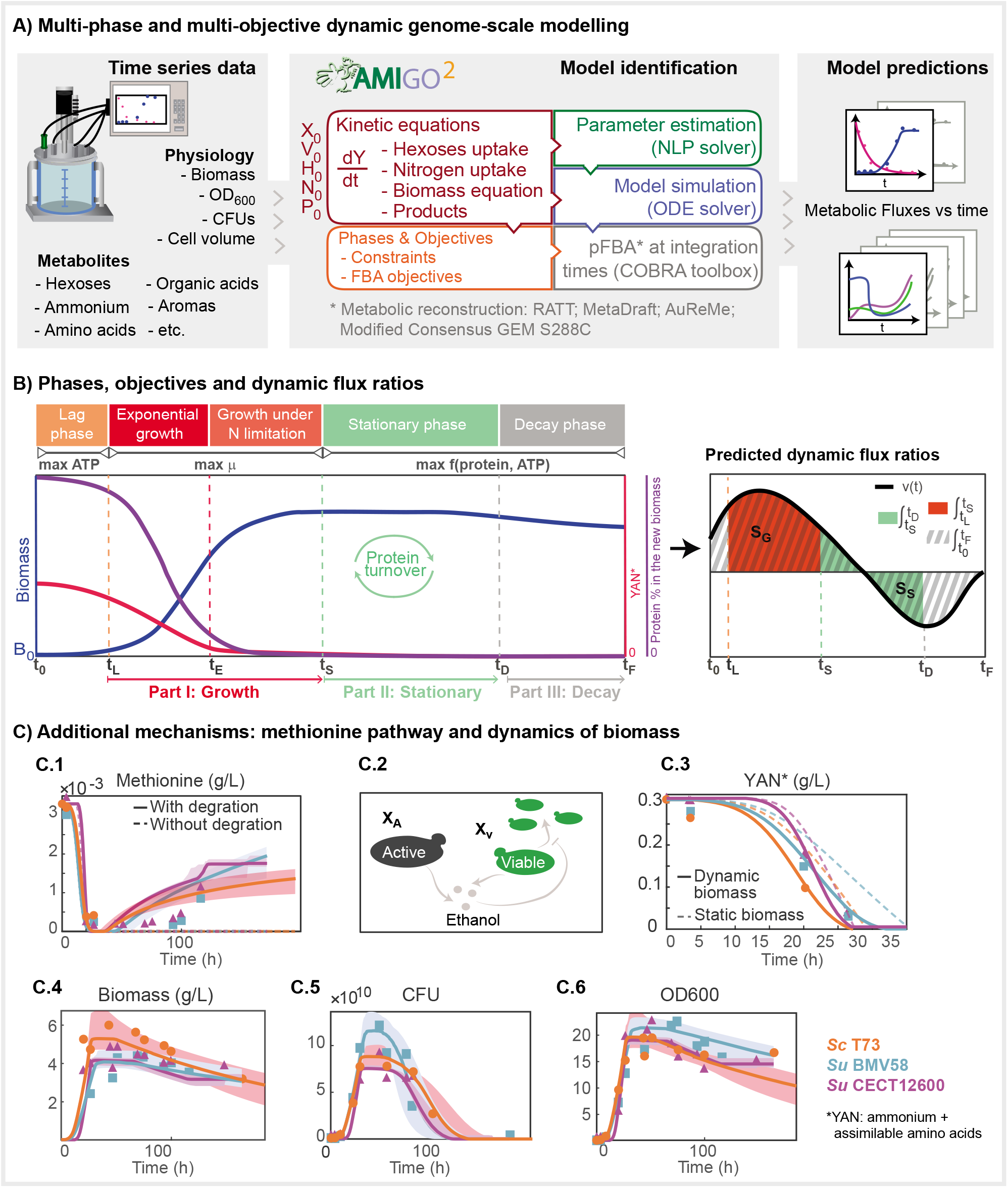
Details on the implementation of the multi-phase and multi-objective dynamic genome-scale model to simulate batch fermentation. A) Implementation, model formulation and solution approach; B) Multi-phase and multi-objective dynamic FBA and methodology to compute dynamic flux rates; the process starts at *t*_0_ = 0 and ends at *t_F_*, the timing of each phase *t_L_, t_E_, t_S_* and *t_D_* is computed through parameter estimation; C) Model improvements through additional mechanisms: C1) model prediction vs the experimental dynamics of methionine with and without its degradation pathway, C2) schematic view of active vs viable biomass in the model, C3) YAN consumption prediction with static and dynamic biomass equations,C4-C6) model predictions vs biomass, CFU and OD600 measurements.

Once inoculated, cells encounter new nutrients and undergo a temporary period of non-replication, the lag-phase, during which we assumed that ATP production is maximized. The exponential growth phase covers only the first hours until nitrogen exhaustion. In this phase, cells maximize growth. During growth under nitrogen limitation cells, still maximizing growth, accumulate carbohydrates. Thenceforward, a substantial fraction of the sugar is consumed during the stationary and decay phases by quiescent cells, which adjust their metabolism to cope with environmental fluctuations. In the latter two phases, we assumed cells maximize both ATP and protein production. Additionally, constraints differ in the different phases.

The general formulation of the FBA problem reads as follows:

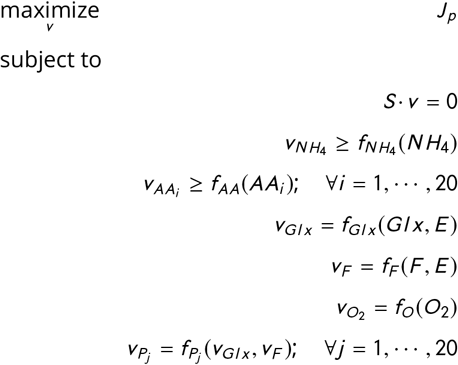

where *J_p_* is the function to be maximized in each phase *p, S* is the stoichiometric matrix, *v* is the vector of fluxes in *mmol*/(*gDWh*), *v_Glx_* and *v_F_* are the fluxes of glucose and fructose, *v_O_2__* is the flux of *O_2_* present only at the beginning of the fermentation, *v_NH_4__* is the flux of ammonium, *v_AAi_* is the exchange rate of the amino acid *i* (covering all 20 amino acids), *v_P_j__* are the constraints associated with the *j* = 1,…, 20 fermentation products considered. *Glx*, *F*, *NH_4_, AA_i_, P_j_* correspond to the concentrations of glucose, fructose, ammonium, amino acids and products, all expressed in (*mmol*/*L*). The Table S2 presents the specific formulation for each phase.

The uptake of glucose and fructose was modelled using Michaelis-Menten (MM) type kinetics with competitive ethanol inhibition (27):

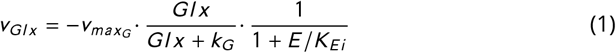

where *v_max_G__* is the maximum uptake rate, *k_G_* is the MM constant, *K_Ei_* is the strength of ethanol inhibitory effect and *E* its concentration (mmol/L). A similar expression *v_F_* exists for fructose (*F*). Additionally, in our case studies, the media was supplemented with sucrose; thus, we included a mass action type expression, characterised by the kinetic constant *k_hydro_*, describing its hydrolysis.

Certain amount of dissolved oxygen is present in the media and consumed during the lag phase (see Figure 2). Its uptake follows:

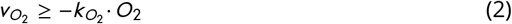

where *v_O_2__* and *k_O_2__* are the oxygen uptake and transport rate constants and *O_2_* the concentration of oxygen in the media.

The uptake of ammonium was modeled by:

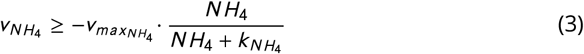

where *NH_4_* is the extracellular concentration of ammonia (mmol/L), *v_maxNH_4__* is the maximum uptake rate achieved, *k_NH_4__* is the MM constant. To avoid an excessive number of parameters, amino acid transport was modeled following mass action kinetics:

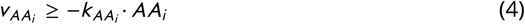

where *AA_i_* is the extracellular concentration of the amino acid (mmol/L) and *k_AA_i__* is the associated kinetic parameter.

**FIG 2.**
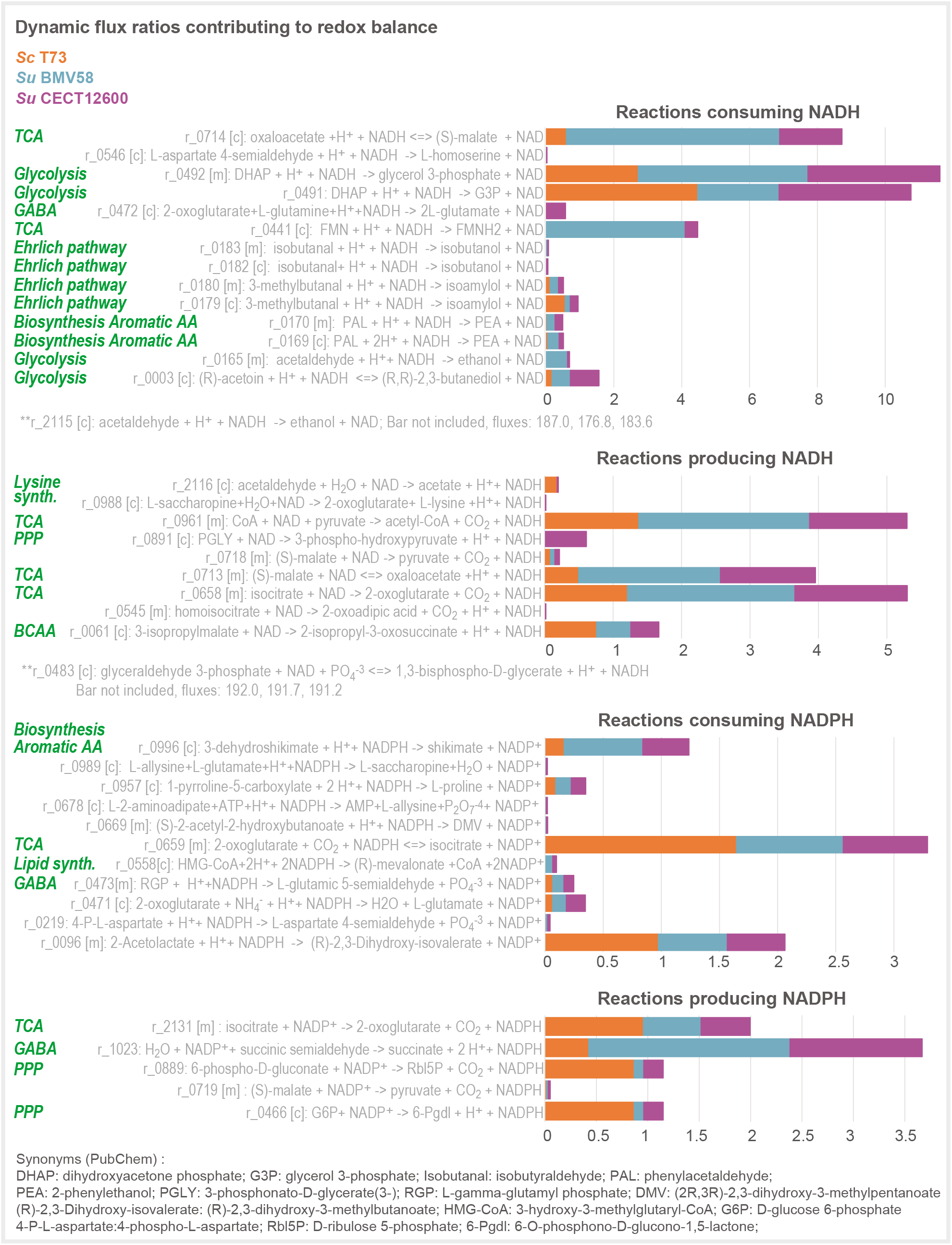
Comparative study of the fluxes through the reactions consuming and producing NADH and NADPH at the stationary phase. Figure illustrates how strains achieve redox balance. The most siginificant differences between strains are found at the level of the TCA cycle, the GABA shunt, the pentose phosphate pathway, the biosynthesis of aromatic amino acids which eventually lead to produce 2-phenylethanol (PEA) and the Ehrlich pathway toward producing isamylol. It is also important to note that *S. uvarum* diverts flux to the production of mevalonate.

Production of alcohols and higher alcohols, carboxylic acids and esters follows mass action kinetics:

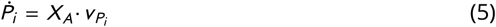

with *X_A_* the active biomass, and the flux *v_P_i__* proportional to the amount of transported hexoses:

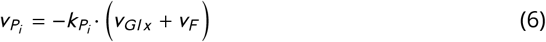

where *P_i_* refers to the excreted product *i* = 1, 20 and *k_P_i__* the production rates.

An exception to this was the formulation of the dynamics of acetate. This metabolite is produced during exponential growth and consumed during stationary phase following mass action kinetics.

### The model of protein turnover

Since nitrogen sources are depleted before the stationary phase, we developed a new model of nitrogen homeostasis that considered protein turnover. The proposed model describes the combined use of the Ehrlich and *de novo* synthesis pathways during stationary and decay phases to guarantee optimal adaptation to perturbations in nitrogen homeostasis.

To introduce protein turnover, we simulated the degradation of the existing protein fraction inside biomass (*Prot*), into a pool of amino acids that subsequently produce new proteins. During stationary and decay phases, the lower bounds on the amino acid uptake are set as:

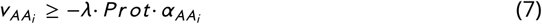

where *λ* is the turnover rate, *Prot* is the concentration of protein and *α_AA_i__* is associated with the stoichiometric coefficient of the amino acid *i* in the protein pseudo reaction. Mathematically, the dynamics of protein content reads:

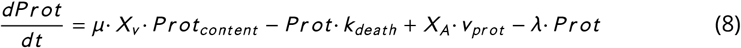

where *μ* is the growth rate, *Prot_content_* is the fraction of protein in the newly formed biomass, *k_death_*(*h*^−1^) is the rate of biomass degradation during decay phase, *X_V_* is the simulated viable biomass (g/L), *X_A_* the simulated active biomass (g/L), *v_Prot_* the protein production rate (*g · gDW*^−1^ · *h*^−1^) and *λ* the protein turnover rate (*h*^−1^).

The degraded proteins (*λ · Prot*) are converted into extracellular amino acids whose concentrations are represented by the following equations:

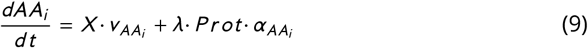

The rates *v_AA_i__* are computed by maximizing protein production (*v_Prot_*) and ATP (*V_ATP_*) while solving the FBA problem:

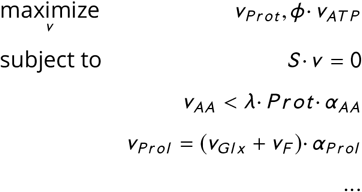

where *ϕ* is estimated for each strain, *S* is the stoichiometric matrix, *v* is the vector of fluxes, *v_AA_i__* is the exchange rate of amino acid *AA_i_, v_P_i__* constraints associated with fermentation products and *v_Prol_* (r_1904) is the amount of excreted proline. The later amino acid accumulates in the extracellular media throughout the fermentation (see Figure 2), likely as a consequence of stored arginine consumption in anaerobic conditions (28). The extracellular dynamics of proline is described as follows:

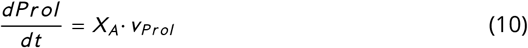

Depending on the kinetic constraints, amino acids can be directly incorporated into proteins or degraded to recover nitrogen for protein production. We observed that those amino acids which lacked pathways for their catabolism or elimination, accumulated in the extracellular compartment. As an example, Figure 1.C1 shows how the model can successfully recover the dynamics of methionine, suggesting the pathway for its degradation – included in our novel reconstruction – could be active during the stationary phase.

### The dynamic biomass equation

A static biomass equation was not able to explain nitrogen assimilation (Figure 1.C2). Therefore we implemented the following dynamic biomass equation (15):

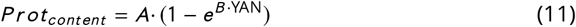

where *A* and *B* are estimated parameters and YAN accounts for the ammonium and free amino acids present in the medium, excluding proline, which is not catabolized under anaerobic conditions.

Furthermore, we assumed that mRNA level was proportional to the protein content (*mRNA = prot/RNA_to_Protein_Ratio*). In this framework, carbohydrates compensate for the variation in protein and mRNA content. Growth-associated ATP maintenance (*GAM*) was also updated to account for the polymerization costs of the different macromolecules (protein, RNA, DNA and carbohydrates):

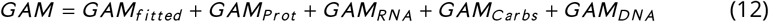

where *GAM_fitted_* is a species or strain-dependent parameter estimated from data and the rest are polymerization costs of the different biomass precursors (adapted from (18)). Additionally, to represent the premature end of fermentations during the decay phase (observed in SuCECT12600), we estimated the non-growth-associated maintenance (NGAM).

In addition, we discriminated between active-able to ferment- and viable cells – able to divide and ferment-to capture the dynamics of CFUs and biomass (Figure 1.C2). The dynamics of active cell mass is represented by the equation:

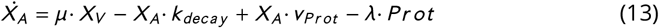

where *X_A_* is the active cell mass (g/L), *μ* is the growth rate computed with pFBA, *k_decay_* is the decay rate (only active during decay), *v_Prot_* (r_4047) is the exchange flux for protein production and *λ* is the turnover rate (both, only active during stationary and decay phases). The behaviour of viable cell mass differed from that of active cell mass by a decline induced by ethanol (29):

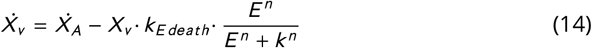

where *X_v_* (g/L) is the growth rate obtained with the constraint based model, *E* is the ethanol concentration (mmol/L) and *k_Edeath_*, *n* and *k* are the parameters controlling susceptibility to ethanol.

The former mechanisms, coupled to parameter estimation, allowed us to predict nitrogen consumption, CFU and biomass dynamics accurately (Figures 1.C3-C6).

### Goodness-of-fit of the model in the case studies

The final model consisted of 46 ordinary differential equations depending on 66 parameters which we estimated from time-series data for all measured external metabolites and biomass. The mean standard deviation on the parameters ranges from 2.5% for SuCECT12600 and a 12.6% for SuBMV58. The reasonably low distribution on the parameters resulted in a reasonably low uncertainty associated with the model simulations (as seen in Figures 2–3).

**FIG 3.**
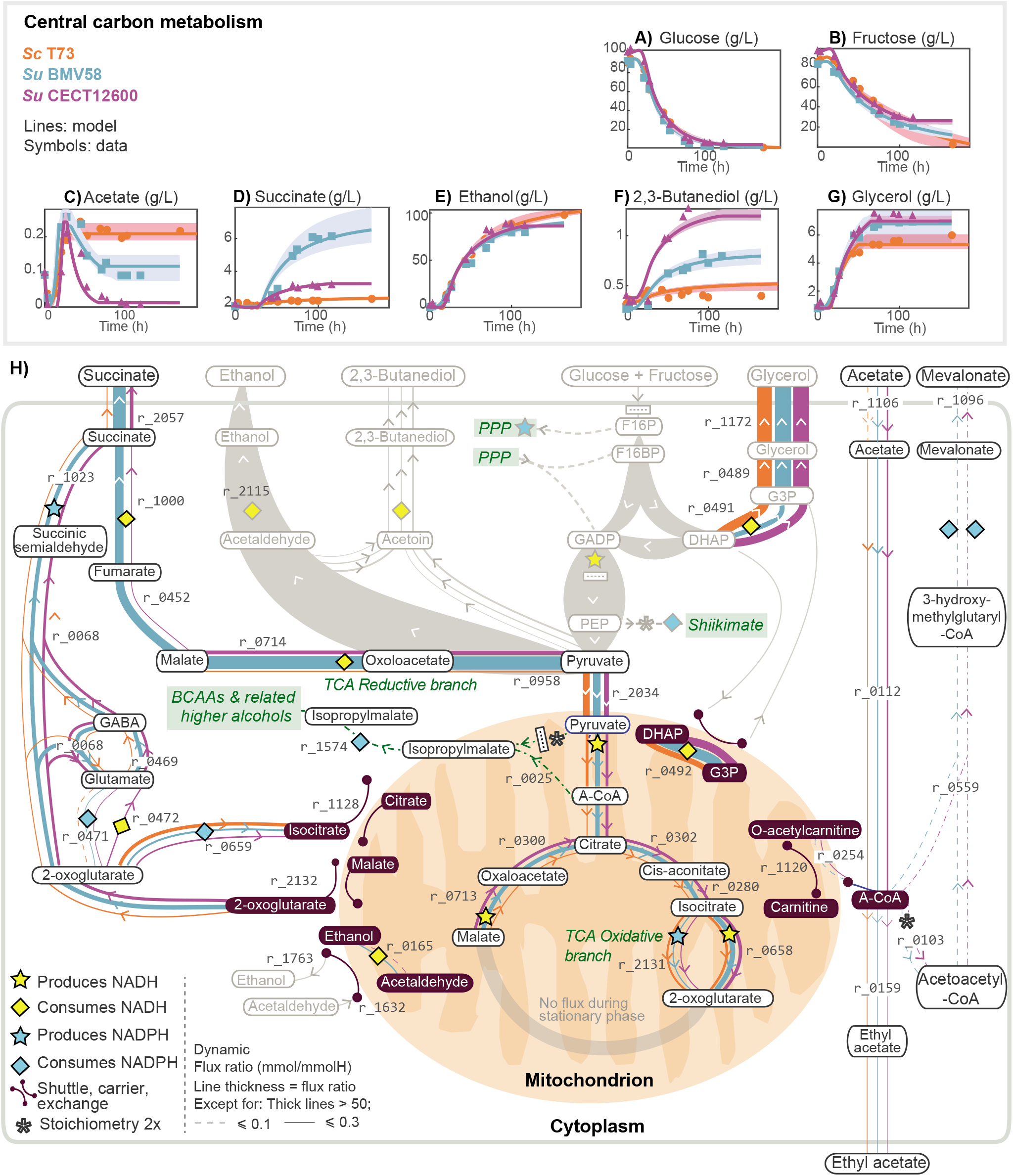
Redox balance in Central Carbon Metabolism. Figures A) to F) show model predictions versus the experimental data extracellular metabolite concentrations associated with glycolysis and central carbon metabolism for the three strains. Figure H) presents the predicted intracellular dynamic flux ratios during the stationary phase, showing how *S. uvarum* and *S. cerevisiae* strains use different redox balance strategies. These differences result in the differential production of relevant external metabolites such as acetate (C), succinate (D), ethanol (E), 2,3-butanediol (F) or glycerol (G). Explicit differences in the pathways in gray are presented in Text S3; otherwise, as indicated in the legend, width of the lines is proportional to the dynamic flux ratio.

The model described the dynamics of our illustrative examples successfully. The best fit to the data plus the associated uncertainty, as computed by the bootstrap, are shown in Text S2 and Figures 2 and 3. We determined the R-squared measure of goodness of fit (*R*^2^) for each measured variable and each strain-based fermentation. The median of the *R*^2^ values are above 0.94 for all strains.

Interested readers may find further details on the parameter estimation in Text 2. Optimal parameter values and *R*^2^ values are reported in Table S3.

### Species behaviour differs significantly in the stationary phase

At the extra-cellular level, the most stricking differences between strains occur in the dynamics of acetate and the yield of succinate (Figures 3C, 3D) and in the production of higher alcohols 2-phenylethanol and isoamyl alcohol (Figures 4B, 4D). We used the model to decipher the metabolic strategies used by the different strains that lead to such differences. The Table S4 reports the dynamic flux ratios as mmol of produced compound per mmol of consumed hexose x 100 (from now on, mmol/mmolH) for those reactions in which the maximum flux ratio value over the three species is above 0.01 mmol/mmolH. Our model predicted that strains use different redox balancing strategies during the stationary phase at the level of central carbon metabolism and the production of higher alcohols. Figure 2, presents a comparison of the dynamic flux ratios for those reactions contributing to redox balance. Interestingly *S. uvarum* strains use the GABA shunt as a NADPH source and acetate to produce mevalonate while the production of 2-phenylethanol is also a pivotal contributor in their cellular redox state.

**FIG 4.**
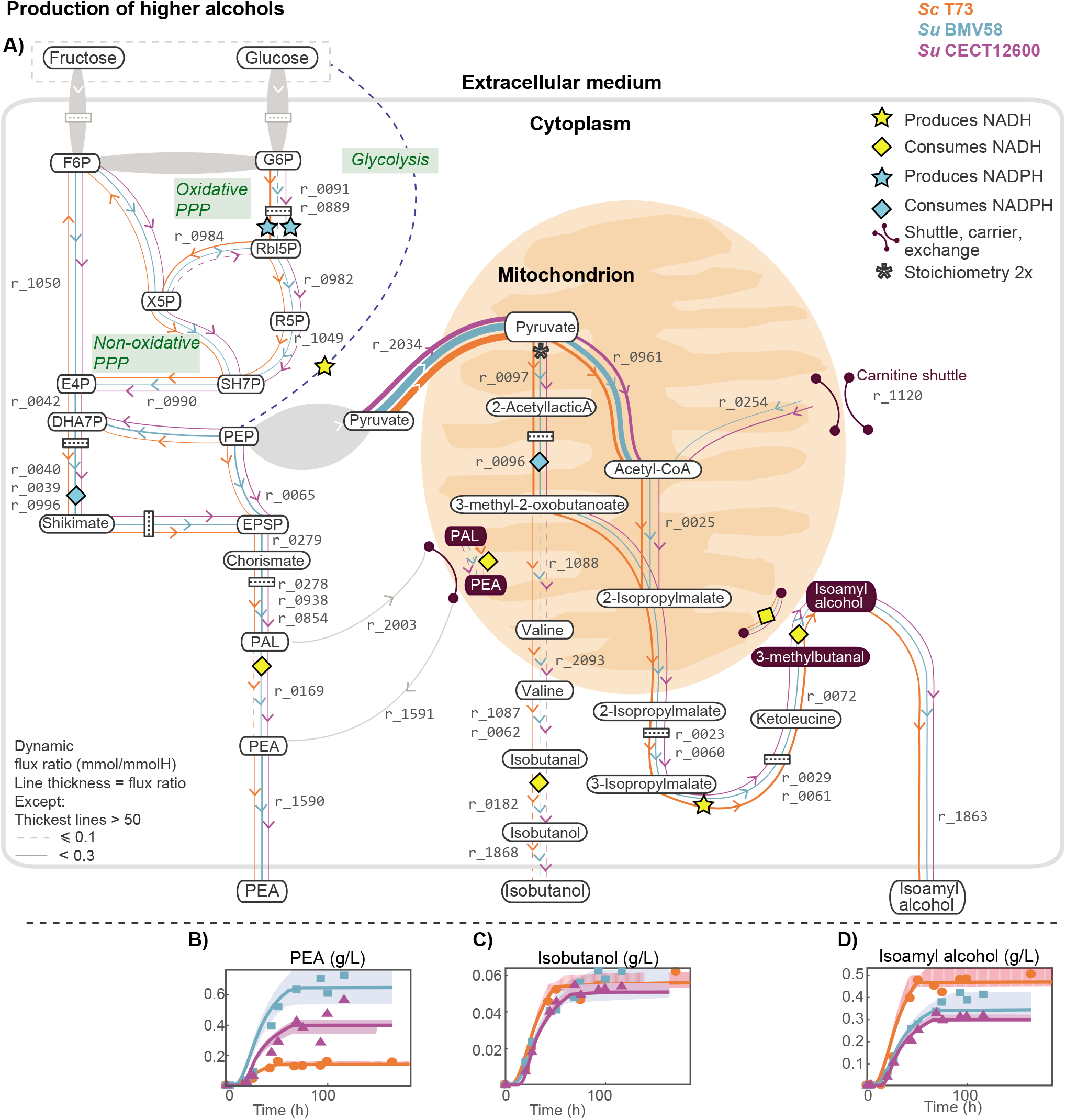
Redox balance in higher alcohol production: Figure A) shows the predicted intracellular flux ratios (above 0.01 mmol/mmolH) related to higher alcohols 2-phenylethanol (PEA), isobutanol and isoamyl alcohol during the stationary phase and their corresponding impact on the redox co-factors balance NADPH/NADP+ and NADH/NAD+. Figures B-D correspond to the comparison between model predictions and raw measures of PEA, isobutanol and isoamyl alcohol, respectively. Other higher alcohols such as methionol and tyrosol, seemed to accumulate in minimal quantities (flux ratio *x* 100 ≃ 0.001) in response to perturbations in the amino acid pool. Remarkably, the flux ratios corresponding to the degradation of amino acids are well below 0.01.

### The GABA shunt as an NADPH source in cryotolerant species

Cells produced the most significant fraction of succinate during the stationary and decay phases, with a significantly higher dynamic flux ratio by SuBMV58 and SuCECT1600 (6.08 and 1.66) than ScT73 (0.42, *r*_2057).

During the decay phase, most succinate was produced through the TCA cycle reductive branch in the two species. However, during the stationary phase, succinate production was distributed between the GABA shunt (ScT73: 0.42, SuBMV58: 2.00, SuCECT12600: 1.31; *r*_1023, Figure 3.H) and the reductive branch of TCA (ScT73: 0.00, SuBMV58: 4.10 and SuCECT12600: 0.35; *r*_1000, Figure 3.H). Remarkably, the GABA shunt was the primordial source of succinate during the stationary phase in ScT73 and SuCECT12600 strains-between 3 and 4.6 times higher than *S. cerevisiae* (Figure 3.H, *r*_0068, *r*_1023). This result suggested an important role of the GABA shunt in the maintenance of the cellular redox state. Incidentally, revisiting data from López-Malo et al. (30) we found a considerable accumulation of GABA (93.59 fold-change) by SuCECT12600 (Table S4).

### Intracellular mevalonate as a reducing equivalent in cryotolerant yeast species

The three strains produced acetate during the growth phase (ScT73: 1.079, SuBMV58: 1.145, SuCECT12600: 1.422; Table S4, *r*_1106) and until the entry into the stationary phase. Afterward, while extracellular acetate concentration remained constant in ScT73, a decrease was observed in both *S. uvarum* fermentations, indicating acetate consumption. As shown in Figure 3.C, our model successfully described these phenotypes.

According to the modeling constraints, the most parsimonious explanation for this observation would have been an operative glyoxylate cycle. However, based on the repression by glucose of the key enzymes of the glyoxylate cycle (i.e. *ICL1* and *MLS1*) and previous intracellular data (30), we decided to block this cycle and explore an alternative hypothesis. As a result, the model suggested that *S. uvarum* strains incorporated the acetate derivative, acetyl-CoA, into mevalonate (SuBMV58: 0.19 and SuCECT12600: 0.10; *r*_0559, *r*_0103) also consuming NADPH. In addition, mevalonate is a reducing equivalent that can be further metabolized into, for example, ergosterol (*r*_0127 in our reconstruction) possibly acting as storage of NADPH in cryotolerant species.

*S. uvarum* strains also used the carnitine shuttle system to transport acetyl-CoA into the mitochondria (SuBMV58: 0.09 and SuCECT12600: 0.38; Figure 3.H, *r*_0254). Inside the mitochondria, acetyl-CoA was used to form isopropylmalate (Figure 4.A) – a precursor of leucine and isoamyl alcohol – or in the TCA oxidative branch towards the synthesis of 2-oxoglutarate (Figure 3.H).

### The production of higher alcohols contributed to the redox balance

Higher alco-hol production was most prominent during the stationary phase for the three strains. Our model predicted that carbon skeletons of isoamyl alcohol, 2-phenylethanol (PEA) and isobutanol were in great part synthesized *de novo* from glycolytic and pentose phosphate pathway intermediates, rather than coming from the catabolism of precursor amino acids (leucine, valine and phenylalanine respectively) (Figure 4.A).

*S. uvarum* strains produced more PEA than ScT73 strain (ScT73: 0.158, SuBMV58: 0.659, SuCECT12600: 0.383; *r*_1590) while the opposite occurs for isoamyl alcohol (ScT73: 0.736,SuBMV58: 0.483, SuCECT12600: 0.394; *r*_1863). We found that the production of PEA and isoamyl alochol contributed substantially to the redox metabolism related to glycerol accumulation. Approximately 43%, 36% and 27% of the glycerol produced by the ScT73, SuBMV58 and SuCECT12600 strains, was attributable to NADH derived from isoamyl alcohol and PEA.

The higher production of PEA observed in *S. uvarum* strains is due to the higher flux through the shikimate pathway (Figure 4.A, *r*_0996, *r*_0279). Interestingly, while ScT73 had a larger flux ratio through the oxidative pentose phosphate pathway (PPP, Figure 4.A, *r*_0091, *r*_0889) partly redirected toward glycolysis, *S. uvarum* simulations reflected the inverse pattern (Figure 4.A, *r*_0984), with glycolytic flux being shifted towards the non-oxidative PPP.

Pyruvate in the mitochondrion showed two different fates: acetyl-CoA (*r*_0961) and 2-acetyllactic acid (*r*_0097). Noticeably, *S. uvarum* strains also contributed to acetyl-CoA using the carnitine shuttle (*r*_0254). 2-acetyllactic acid can further be converted to 3-methyl-2-oxobutanoate, consuming one NADPH (*r*_0096), which also showed two different fates. The production of 2-isopropylmalate (*r*_0025), leading to isoamyl alcohol (*r*_0072, *r*_0179); or 3-methyl-2-oxobutanoate, leading to the synthesis of isobutanol via the Ehrlich pathway (*r*_1087, *r*_0062, *r*_0182). All strains showed similar productions of isobutanol (ScT73: 0.1, SuBMV58: 0.084, SuCECT12600: 0.078; r_1868).

Isoamyl alcohol and 2-phenylethanol significantly impacted redox metabolism, mainly when biosynthetic requirements were low. Despite that the NADH requirements of 2-phenylethanol are four times smaller than those of isoamyl alcohol, the former also impacted the NADP+/NADPH ratio. The PPP branch followed to produce 2-phenylethanol precursor erythrose-4 phosphate might result in NADPH or NADP+ accumulation. We predicted that, during the stationary phase, 2-phenylethanol produced through the chorismate synthesis pathway (downstream of the non-oxidative PPP), provided some of the excess NADP+ required to transform succinic semialdehyde into succinate.

## DISCUSSION

Genome scale models have the potential to decipher how non-conventional yeast species use metabolism to produce industrially relevant products and tolerate specific stressors, such as cold temperatures. This study aimed to develop a dynamic genomescale model to investigate the dynamics of yeasts primary and secondary metabolism in batch cultures. To generate biological hypotheses for modeling, we considered the description of the metabolism of *S. cerevisiae* and cryotolerant *S. uvarum* strains in a rich medium (grape must) fermentation.

The first question in this research was how to model all phases in the batch process: lag, exponential growth, limited nitrogen growth, stationary and decay. Prior studies focused on the exponential growth phase and were based on available reconstructions with many missing reactions, particularly those related to secondary metabolism (14, 15, 31, 32). Also, their static nature hinders the description of the sequential nature of amino acid consumption (33).

As a first step, we needed to extend a yeast genome-scale reconstruction to account for the production of higher alcohols, carboxylic acids or esters. We extended the Yeast8 consensus model incorporating missing reactions and metabolites. A similar curation process has been recently applied to the iMM904 reconstruction (34). The authors fitted the model to data from the literature concluding that further curation and adaptations were necessary to successfully predict metabolism and biomass dynamics. We experienced such difficulties in our first iterations in the modeling process and introduced several new features to obtain more accurate simulations of carbon and nitrogen metabolism throughout time.

A critical aspect for an improved accuracy was the multi-phase multi-objective dynamic FBA scheme. Previous works focused on ATP consumption to explain the metabolism after depletion of the limiting nutrient (35, 36, 15), we incorporated the production of protein as a cellular objective (together with protein degradation) with accurate results. Also, we modeled protein turnover to account for the uptake of amino acids and inorganic nitrogen in the stationary phase. To the best of our knowledge, this is the first dynamic genome-scale metabolic model describing nitrogen homeostasis during the stationary phase. Finally, we introduced a dynamic biomass equation which further improved the model accuracy. This result agrees with observations by previous studies (37, 38), or (39) pointing out the relevance of detailing biomass composition in a context-specific manner.

The second question in this research was to decipher the differences in the metabolism of three strains of two different species, *S. cerevisiae* and *S. uvarum,* in rich medium (grape must) fermentation. Recently, Minebois et al. (40) hypothesized that *S. cerevisiae* and *S. uvarum* species might have different redox balance strategies. Notably, the model confirmed this hypothesis and brought novel insights into the specific routes used by the two species.

Our predictions suggest alternative pathways for cryotolerant species to produce succinate and consume acetate. In principle, yeasts might form succinate via four main pathways, all based on the reactions of the TCA cycle (41). Selected pathway depends on the environmental conditions and strain. Our model predicted that ScT73 and SuBMW58 produced overall the most succinate via the TCA reductive branch, in agreement with Camarasa et al. (42). However, our results also suggest an important role of the TCA oxidative branch until 2-oxoglutarate for the *S. uvarum* strains during the stationary phase. This result is consistent with the recent intracellular data obtained by (25) who observed a noticeable intracellular accumulation of 2-oxoglutarate in SuBMV58.

One somewhat unexpected finding of the model was the extent to which the GABA shunt would contribute to succinate formation during the stationary phase, an effect particularly evident in the case of *S. uvarum* strains. The role of this pathway is not fully understood in yeast (43, 44, 45). Bach et al. (43) observed that glutamate decarboxylase (*GAD1*) was poorly expressed when succinate was produced in *S. cerevisiae* and that the GABA shunt played a minor role in redox metabolism. On the contrary, Coleman et al. (46) showed that *GAD1* expression is required for oxidative stress tolerance in *S. cerevisiae.* Similarly, Cao et al. (44) showed that *GAD1* confers resistance to heat stress effect that migth be related to NADPH production. Additionally, *GAD1* was up-regulated during the stationary phase under nitrogen starvation (47, 43, 45); and (30) observed high intracellular GABA levels in cryotolerant species *S. uvarum.*

Recently, Liu et al. (48) found clear indication that the GABA shunt may be involved in supplying NADPH for lipid synthesis in the oleaginous yeast *Yarrowia lipolytica.* Also, Bach et al. (43) showed that *S. cerevisiae* can degrade GABA into succinate or *γ*-Hydroxybutyric acid (GHB) and that GHB was used to form the polymer polyhydroxybutyrate (PHB). Also noteworthy is that PHBs are synthesized by numerous bacteria as carbon and energy storage compounds (49). PHBs are also strongly associated with bacterial cold tolerance (50) suggesting a similar function in yeast.

Another important finding is that *S. uvarum* strains consume acetate once nitrogen sources are depleted, coinciding with the extracellular accumulation of succinate. This finding was also reported by Kelly et al. (51), who showed that a *S. uvarum* yeast isolate can metabolize acetate to significantly lower acetic acid, ethyl acetate, and acetaldehyde in wine. The model predicted that some of the acetate carbon was directed toward mevalonate, which is in line with recent experimental work by (25). According to Bach et al. (43), a route for acetyl-CoA incorporation into PHB polyester through 3-hydroxybutyrate-CoA seems plausible. However, this hypothesis is not taken into account by the genome-scale reconstructions.

The fact that López-Malo et al. (30) found high intracellular GABA and GHB in cryotolerant strains grown in synthetic must (without GABA) at low-temperature, and the flux predicted in the present work, suggest that *S. uvarum* stores lipids or polyesters (i.e., PHBs) as reducing equivalents to withstand oxidative stress induced by low temperatures. The role of the GABA shunt and the production of reducing equivalents in the metabolism of cryotolerant species, may be plausible routes worth exploring.

Our model also predicted that the carbon skeletons of higher alcohols (e.g., isobutanol and isoamyl alcohol) were mainly synthesized *de novo* rather than from the incorporation and catabolism of amino acids (e.g., leucine and valine). This result agrees with the findings of Crépin et al. (52) who explored the fate of the carbon backbones of aroma-related exogenous amino acids using 13C isotopic tracer experiments. Similarly, our results indicate that 2-phenylethanol was mostly synthesized *de novo.* We hypothesize that a positive contribution in glycerol content may also explain why the production of 2-phenylethanol and isoamyl alcohol is a conserved evolutionary trait in yeasts.

Interestingly, the model predicts that most other higher alcohols (tyrosol, methionol, etc.) accumulate in small amounts due to perturbations in the amino acid pool. These results confirm the hypothesis raised by Shopska et al. (53) who suggested that the two schemes to produce higher alcohols – Ehrlich and *de novo* synthesis-are not in contradiction but two extremes of a common mechanism. Incidentally, Yuan et al. (54) showed that the assembled leucine biosynthetic pathway coupled with the Ehrlich degradation pathway results in high-level production of isoamyl alcohol. The fact that model predictions are consistent with numerous previous findings lead us to conclude that the present model (along with the provided code) can simulate yeast metabolism in batch culture in a general chemically characterized medium; the only requirement would be to update the metabolic reconstruction if required for the specific yeast species. The model can also be used to explore and engineering novel metabolic pathways towards specific bioproducts.

## MATERIALS AND METHODS

### Yeast strains

In this study, three yeast strains belonging to *S. cerevisiae* and *S. uvarum* species were used: the commercial strain, T73 (Lalvin T73 from Lallemand Montreal, Canada), originally isolated from wine in Alicante, Spain (55) was selected as our wine *S. cerevisiae* (ScT73) representative; the commercial strain BMV58 (SuBMV58, Velluto BMV58 from Lallemand Montreal, Canada), originally isolated from wine in Utiel-Requena (Spain) and the non-commercial CECT12600 strain, isolated from a non-fermentative environment (SuCECT12600, Alicante, Spain).

### Fermentation experiments

Fermentation assays were performed with grape must obtained from the Merseguera white grapes, collected in the 2015 vintage in Titaguas (Spain) and stored in several small frozen volumes (4 l, −20°C). Before its use, the must was clarified by sedimentation for 24 h at 4°C and sterilized by adding dimethyl dicarbonate at 1 ml.l^−1^. All fermentations were performed in 3*x* independent biological replicates in 500 ml controlled bioreactors (MiniBio, Applikon, the Netherlands) filled with 470 ml of natural grape must. Each bioreactor was inoculated using an overnight starter culture cultivated in Erlenmeyer flasks containing 25 ml of YPD medium (2% glucose, 0.5% peptone, 0.5% yeast extract) at 25°C, 120 rpm in an agitated incubator (Selecta, Barcelona, Spain). Strain inoculation was done at OD600=0.100. The dynamics of the fermentation was registered using different probes and detectors to control and measure temperature, pH, dissolved oxygen (Applikon, The Netherlands) and effluent carbon dioxide level (INNOVA 1316 Multi-Gas Monitors, LumaSense Technologies). Data were integrated into the BioExpert software tools (Applikon, The Netherlands). The fermentation was complete when a constant sugar content was reached as measured by HPLC.

### Sampling and quantification of extracellular metabolites

Extracellular metabolites, including sugars, organic acids, main fermentative by-products, and yeast assimilable nitrogen (YAN) were determined at ten sampling times during the fermentation. Residual sugars (glucose, fructose), organic acids (acetate, succinate, citrate, malate and tartrate) and the main fermentative by-products (ethanol, glycerol and 2.3 butanediol) were quantified using HPLC (Thermo Fisher Scientific, Waltham, MA) coupled with refraction index and UV/VIS (210 nm) detectors. Metabolites were separated through a HyperREZ XP Carbohydrate H+ 8 *μ*m column coupled with a HyperREZ XP Carbohydrate Guard (Thermo Fisher Scientific, Waltham, MA). The analysis conditions were: eluent, 1.5 mM of H2SO4; 0.6 ml.min^−1^ flux and a 50°C oven temperature. For sucrose determination, the same HPLC was equipped with a Hi-Plex Pb, 300 x 7.7 mm column (Agilent Technologies, CA, USA) and the following analysis conditions were used: eluent, Milli-Q water; 0.6 ml.min^−1^ flux and oven temperature of 50°C. The retention times of the eluted peaks were compared to those of commercial analytical standards (Sigma-Aldrich, Madrid, Spain). Metabolite concentrations were quantified by the calibration graphs (R2 value > 0.99) of the previously obtained standards from a linear curve fit of the peak areas using ten standard mixtures.

Determination of yeast assimilable nitrogen in the form of amino-acids and ammonia was carried out following the same protocol as (56). A volume of supernatant was removed from the fermenter, and amino acids and ammonia separated by UPLC (Dionex Ultimate 3000, Thermo Fisher Scientific, Waltham, MA) equipped with a Kinetex 2.6u C18 100A column (Phenomenex, Torrance, CA, USA) and Accucore C18 10 x 4.6 mm 2.6 *μ*m Defender guards (Thermo Fisher Scientific, Waltham, MA). For derivatization, 400 *μ*l of the sample was mixed with 430 *μ*l borate buffer (1M, pH 10.2), 300 *μ*l absolute methanol and 12 *μ*l of diethyl ethoxymethylenemalonate (DEEMM), and ultra-sonicated for 30 min at 20°C. The ultra-sonicated sample was incubated up at 80°C for 2 hours to allow the complete degradation of excess DEEMM. Once the derivatization finished, the sample was filtered with 0.22 *μ*m filter before injection. The target compounds in the sample were then identified and quantified according to the retention times, UV-vis spectral characteristics and calibration curves (R2 value > 0.99) of the derivatives of the corresponding standards. Amino acid standard (ref. AAS18), asparagine and glutamine purchased from Sigma-Aldrich were used for calibration.

### Higher alcohols and esters

We also determined the concentrations of higher alcohols and esters for each sampling time. Volatile compound extraction and gas chromatography were performed following the protocol of Rojas et al. (2001). Extraction was performed using headspace solid phase-micro-extraction sampling (SPME) with poly-dimethylsiloxane (PDMS) fibers (Supelco, Sigma-Aldrich, Barcelona, Spain). Aroma compounds were separated by GC in a Thermo TRACE GC ULTRA chromatograph (Thermo Fisher Scientific, Waltham, MA) equipped with a flame ionization detector (FID), using a HP-INNOWAX 30 m x 0.25 mm capillary column coated with a 0.25 mm layer of cross-linked polyethylene glycol (Agilent Technologies, CA, USA). Helium was the carrier gas used (flow 1 ml.min^−1^). The oven temperature program was: 5 min at 60°C, 5°C.min^−1^ to 190°C, 20°C.min^−1^ to 250°C and 2 min at 250°C. The detector temperature was 280°C, and the injector temperature was 220°C under splitless conditions. The internal standard was 2-heptanone (0.05% w/v). Volatile compounds were identified by the retention time for reference compounds. The quantification of the volatile compounds was determined using the calibration graphs of the corresponding standard volatile compounds.

### Physiological and biomass parameters

Physiological and biomass parameters, including OD600, dry weight (DW), colony-forming units (CFUs) and average cell diameter (ACD), were determined at each sample time, providing that the cell sample was sufficient to perform the corresponding measure. DW determination was performed by centrifuging 2 ml of the fresh sample placed in a pre-weighed Eppendorf tube in a MiniSpin centrifuge (Eppendorf, Spain) at maximum speed (13.200 rpm) for 3 min. After centrifugation, the supernatant was carefully removed, the pellet washed with 70% (v/V) ethanol, and centrifuged in the same conditions. After washing, the aqueous supernatant was removed carefully and the tube placed in a 65°C oven for 72h. DW was finally obtained by measuring the mass weight difference of the tube with a BP121S analytical balance (Sartorius, Goettingen, Germany). OD600 was measured at each sampling time using a diluted volume of sample and a Biophotometer spectrophotometer (Eppendorf, Germany). CFUs were determined using a 100-200 *μ*l of a diluted volume of samples plated in YPD solid medium (2% glucose, 2% agar, 0.5% peptone, 0.5%yeast extract) and incubated two days at 25°C. The resulting colonies were counted with a Comecta S.A Colony Counter. Only plates with CFUs between 30 and 300 were used to calculate the CFUs of the original sample. For ACD determination, a volume of cell sample was diluted into a phosphate-buffered saline solution and cell diameter measured using a Scepter Handled Automated Cell Counter equipped with a 40 *μ*m sensor (Millipore, Billerica, USA).

### Orthology analysis and genome-scale metabolic reconstruction

Genomes of ScT73, SuBMV58 and SuCECT12600 were sequenced and assembled in previous works ((57); Macías et al., (unpublished)). Genome assemblies were annotated by homology and gene synteny using RATT (58). This approach let us transfer the systematic gene names of *S. cerevisiae* S288c annotation (59) to our assemblies and, therefore, to select only those syntenic orthologous genes in T73, CECT12600 and BMV58 genomes for subsequent analyses.

We added to the consensus genome-scale reconstruction of *Saccharomyces cerevisiae S288C*(*v.8.3.2*) metabolites and reactions related to amino acid degradation and higher-alcohols and esters formation. This refined model was then used as a template for reconstructing strain-specific genome-scale models for SuBMV58, SuCECT12600 and ScT73. MetaDraft, AuReMe and the results from the orthology analysis were used to create the strain-specific models.

### Flux balance analysis

Flux balance analysis (FBA) (60, 61) is a modeling framework based on knowledge of reaction stoichiometry and mass/charge balances. The framework relies on the pseudo steady-state assumption (no intracellular accumulation of metabolites occurs). This is captured by the well known expression:

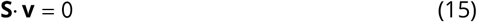

where **S** is stoichiometric matrix of (n metabolites by m reactions) and **v** is a vector of metabolic fluxes. The number of unknown fluxes is higher than the number of equations and thus the system is undetermined. Still it is possible to find a unique solution under the assumption that cell metabolism evolves to pursue a predetermined goal which is defined as the maximisation (or minimisation) of a certain objective function (*J*):

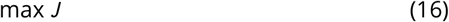

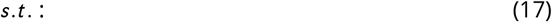

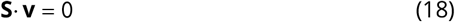

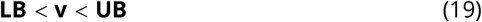

where **LB** and **UB** correspond to the lower and upper bounds on the estimated fluxes. Examples of objective functions *J* include growth rate, ATP, or the negative of nutrient consumption, etc.

Typically, multiple optimal solutions exist for a given FBA problem. In parsimonious FBA (pFBA), the result is the most parsimonious of optimal solutions, i.e., the solution that achieves the specific objective with the minimal use of gene products and the minimization of the total flux load (62).

### Parameter estimation

The aim of parameter estimation is to compute the unknown parameters – growth related constants and kinetic parameters – that minimize some measure of the distance between the data and the model predictions. The maximum-likelihood principle yields an appropriate measure of such distance (63):

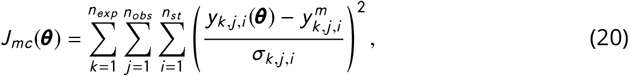

where *n_exp_, n_obs_* and *n_st_* are, respectively, the number of experiments, observables (measured quantities), and sampling times while *σ_k,j,i_* represents the standard deviation of the measured data as obtained from the experimental replicates. 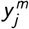 represents each of the measured quantities, *X^m^* and *C^m^* in our case, and *y_j_*(***θ***) corresponds to model predicted values, *X* and *C*. Observation functions were included for *CFUs* and *OD*600 in order to scale viable cell-mass (*X_v_*) and active cell-mass (*X_A_*), respectively.

Parameters are estimated by solving a nonlinear optimization problem where the aim is to find the unknown parameter values (***θ***) to minimize *J_mc_*(***θ***), subject to the system dynamics – the model- and parameter bounds (64).

### Uncertainty analysis

In practice, the value of the parameters ***θ*** compatible with noisy experimental data is not unique, i.e., parameters are affected by some uncertainty (64). The consequence of significant parametric uncertainty is that it may impact the accuracy of model predictions.

To account for model uncertainty, we used an **ensemble approach**. To derive the ensemble, we apply the bootstrap smoothing technique, also known as bootstrap aggregation (the Bagging method) (65, 66). The bagging method is a well established and effective ensemble model/model averaging device that reduces the variability of unstable estimators or classifiers (66). The underlying idea is to consider a family of models with different parameter values **Θ** = [***θ***_1_… ***θ***_*N*_]^*T*^ compatible with the data ***y***^*m*^, when using the model to predict untested experimental setups. The matrix of parameter values **Θ** consistent with the data is obtained using *N* realizations of the data obtained by bootstrap (67). Each data realization has the same size as the complete data-set, but it is constructed by sampling uniformly from all replicates (3 biological replicates per sampling time). Within each iteration, each replicate has an approximate chance of 37% of being left out, while others might appear several times. The family of solutions, **Θ**, is then used to make *N* predictions (dynamic simulations) about a given experimental scenario. The median of the simulated trajectories regards the model prediction, while the distribution of the individual solutions at a given sampling time provides a measure of the uncertainty of the model.

### Analysis of dynamic metabolic fluxes

We selected the most relevant metabolic pathways using a score, which provides a measure of the net flux over time during growth and stationary phases. In particular, we computed the integral of each flux multiplied by the biomass (mmol · h^−1^) over time and normalised its value with the accumulated flux of consumed hexoses (glucose and fructose):

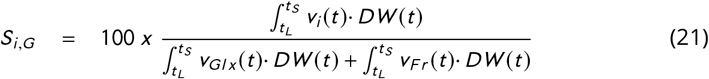

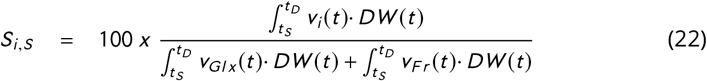

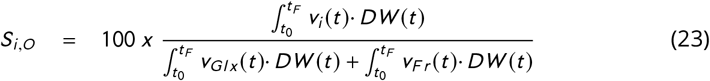

where *S_i,G_* corresponds to the score of the flux *i* during growth, *S_i,s_* corresponds to the score during the stationary, and *S_i,D_* decay phases, *v_i_*(*t*) (mmol · h^−1^ · DW^−1^) is the flux under scrutiny, *v_Glx_*(*t*) (mmol · h^−1^ · DW^−1^) is the flux of glucose, *v_Fr_*(*t*) (mmol · h^−1^ · DW^−1^) is the flux of fructose, and DW is the predicted dry-weight biomass (g). Results correspond to mmol of produced compound per mmol of consumed hexose x 100 (denoted as mmol/mmolH). Score values indicate the overall impact of each reaction in the net oxidation or reduction of electron carriers during the given phase of the fermentation.

### Numerical tools

To automate the modeling pipeline we used the AMIGO2 toolbox (68). To solve the dFBA problem we used a variable-step, variable-order Adams-Bashforth-Moulton method to solve the system of ordinary differential equations that describe the dynamics of the extracellular metabolites. At each time step the pFBA problem was solved using the COBRA Toolbox (69). The global optimiser *Enhanced Scatter Search* (eSS, (70)) was used to find the optimal parameter values in reasonable computational time.

The ensemble model generation procedure is computationally intensive. However, since each parameter estimation instance in the ensemble is an entirely independent task, we were able to solve this problem in less than a day using 60 CPU cores on a Linux cluster. These tasks were automated with the help of bash scripts and the Open Grid Scheduler. All the scripts necessary to reproduce the results are distributed (https://sites.google.com/site/amigo2toolbox/examples.

## SUPPLEMENTAL MATERIAL FILE LIST

- **Table S1. Novel metabolic reconstruction additions.** Table **Sheet 1** presents the list of reactions and metabolites added to the novel metabolic reconstruction. Among the metabolites added, 13 aroma compounds were included, namely, methionol, ethyl-hexanoate, ethyl-octanoate, 1-hexanol, 1-octanol, 4-tyrosol, hexanal, hexyl-acetate, octyl-acetate, benzyl-acetate, 4-hydroxyphenylacetaldehyde and 3-methylsulfanylpropanal. Table **Sheet 2** presents the chemical family (alcohol, ester or aldehyde) and several aroma descriptors for each of the added compounds.
- **Table S2. Detailed mathematical formulation (objective and constraints) of the multi-phase multi-objective pFBA implementation**.
- **Table S3. Optimal parameter values and goodness-of-fit of the model.** Table **Sheet 1** presents the list of all parameters together with their optimal value as achieved by the bootstrap approach for the three individual strains. The comparison between the wine strains ScT73 and SuBMV58, reveals substantial differences – above 100% relative difference, in the growth-associated ATP maintenance; the rate of transport of specific amino acids and hexoses and the production rate of succinate. The natural strain showed that the lag phase is more than three times longer for the natural strain; besides, the non-growth associated maintenance is practically zero for wine strains while it is around 0.8 for the SuCECT12600 strain. The uptake of various amino acids varies significantly between *S. uvarum* strains. In particular, the uptake rate for threonine is 740% higher for SuCECT12600 than for SuBMV58. Table **Sheet 2** presents the R^2^ goodness-of-fit as computed for all measured compounds and biomass for the three individual strains. The vast majority of the coefficients were above 0.9. Some negative values appeared −10 out 141-typically associated with low signal-to-noise ratio and high data dispersion observed in the measured variable, e.g., cysteine.
- **Table S4. Dynamic flux ratios and transcriptomic data. Sheets 1-4** present the overall, growth, decay and stationary dynamic flux ratios respectively mmol of produced compound per mmol of consumed hexose x 100 (from now on, mmol/mmolH). Only those reactions for which the maximum flux ratio value over the three species is above 0.01 mmol/mmolH are reported.textbfSheet 4 presents the (unpublished) data by López-Malo et al. (30) showing substantial accumulation of GABA in *S. uvarum* strains.
- **Text S1. Orthology analysis and genome-scale metabolic reconstruction.** The text further elaborates on the steps followed to obtain individual strains reconstructions.
- **Text S2. Bootstrap parameter estimation of the model.** The text presents further numerical results such as the convergence curves of the bootstrap approach, the distribution of parameter values, and the plots of the model vs data for all measured states not included in the main figures (aminoacids, carbon, nitrogen and O_2_, carboxylic acids, esthers and higher alcohols).
- **Text S3. Detailed description of the redox balance mechanisms used by the different strains.** The text discusses in the detail production of ethanol, glycerol, succinate, 2,3-butanediol, higher alcohols; production and consumption of acetate and contributions to the redox balance.

## ACKNOWLEDGMENTS

This project has received funding from MCIU/AEI/FEDER, UE (grant references: RTI2018-093744-B-C31, RTI2018-093744-B-C32, RTI2018-093744-B-C33 and PID2019-104113RB-I00) and Xunta de Galicia (IN607B 2020/03). RM was supported by an FPI grant from the Ministerio de Economía y Competitividad, Spain (ref. BES-2016-078202). SNM acknowledges funding from CONICYT Becas Chile grant 72180373. SNM and BT acknowledge support from YogurtDesign, EraCoBioTech grant 053.80.733.

## Author contributions

DH implemented the model. RM performed the experiments. SM performed the reconstruction of strain-specific genome-scale models. LGM assembled and annotated the genome. DH, SM and LGM participated in the gap filling process of metabolic functions in the model and in the development of the biomass equations. EBC, RPT, AQ, BT and EB coordinated the work and design of the study/pipeline. DH, RM, SM, LGM, EBC wrote the original manuscript. All authors read, edited, and approved the final paper.

